# Genomic investigation reveals contaminated detergent as the source of an ESBL-producing *Klebsiella michiganensis* outbreak in a neonatal unit

**DOI:** 10.1101/568154

**Authors:** Paul Chapman, Brian M. Forde, Leah W. Roberts, Haakon Bergh, Debra Vesey, Amy V. Jennison, Susan Moss, David L. Paterson, Scott A. Beatson, Patrick N. A. Harris

## Abstract

**Background:** *Klebsiella* species are problematic pathogens in neonatal units and may cause outbreaks, for which sources of transmission can be challenging to elucidate. We describe the use of whole genome sequencing (WGS) to investigate environmental sources of transmission during an outbreak of extended-spectrum-β-lactamase (ESBL)-producing *Klebsiella michiganensis* colonizing neonates.

**Methods:** Ceftriaxone-resistant *Klebsiella* spp. isolated from neonates (or their mothers) and the hospital environment were included. Short-read (Illumina) and long-read (MinION, Oxford Nanopore Technologies) sequencing was used to confirm species taxonomy, define antimicrobial resistance genes and determine phylogenetic relationships using single nucleotide polymorphism (SNP) profiling.

**Results:** A total of 21 organisms (10 patient-derived and 11 environmental isolates) were sequenced. Standard laboratory methods identified the outbreak strain as an ESBL-producing *Klebsiella oxytoca*, but taxonomic assignment from WGS data suggested closer identity to *Klebsiella michiganensis.* Strains isolated from baby bath drains and multiple detergent dispensing bottles were either identical or closely related by SNP comparison. Detergent bottles contaminated by *K. michiganensis* had been used for washing milk-expressing equipment. No new cases were identified once the detergent bottles were removed and the baby baths decommissioned.

**Conclusions:** Environmental reservoirs may be an important source in outbreaks of multi-drug resistant organisms. WGS, in conjunction with traditional epidemiological investigation, can be instrumental in revealing routes of transmission and guiding infection control responses.

**Key points:**

- *K. michiganensis* can be misidentified as *K. oxytoca* and is probably under-recognized as a nosocomial pathogen
- Whole genome sequencing of neonatal and environmental isolates during an outbreak of ESBL-producing *K. michiganensis* confirmed contaminated detergent and sinks to be the source

## Introduction

Outbreaks of ESBL-producing Enterobacterales within neonatal intensive care units are most commonly caused by *Klebsiella* species and may be associated with significant morbidity and mortality (1). *Klebsiella michiganensis* was first identified from a toothbrush holder in a home in Michigan and was initially identified as *Klebsiella oxytoca*, to which it is closely related (99% nucleotide sequence identity in the 16S rRNA gene sequence).(2) Since characterization, *K. michiganensis* has been reported in clinical settings, including as a cause of diarrhea in a hematopoietic stem cell transplant recipient (3), from a sample of fluid from an abdominal fistula (4), in an investigation of hospital-acquired colonization by a New Delhi metallo-β-lactamase-producing organism (5) and as an invasive, *Klebsiella pneumoniae* carbapenemase (KPC)-producing pathogen causing bloodstream infection(6). While the role of *K. oxytoca* in nosocomial outbreaks is established (7–11), there have been no reported outbreaks involving *K. michiganensis* to date.

Similar to *K. oxytoca, K. michiganensis* carries a chromosomally-encoded OXY-type (Ambler class A) β-lactamase (12, 13) (also labelled as K1 in *K. oxytoca* (14)) which mediates resistance to amino- and carboxy-penicillins (e.g. ampicillin, ticarcillin, temocillin). Over-expression of OXY enzymes arising from mutations in regulatory genes (15–17) can lead to phenotypic resistance to multiple β-lactams. Constitutive hyperproducers may also be selected during antibiotic therapy.(18) OXY-type enzymes may give a false-positive result on extended-spectrum β-lactamase (ESBL) confirmation testing by the clavulanate combination disk method (19), yet are not ‘classical’ ESBLs (i.e. Bush-Jacoby Group 2be enzymes (20)).

Outbreaks of multi-drug resistant (MDR) organisms in neonatal units are of significant concern as empirical antibiotic strategies defined in most guidelines may have limited efficacy against MDR gram-negative bacilli. Hospitalized newborns, especially when premature, are at risk for infection caused by *Klebsiella* spp.(21), which account for a significant proportion of nosocomially-acquired neonatal infections, particularly in developing countries.(22)

We describe an outbreak of ESBL-producing *K. michiganensis* (initially misidentified as *K. oxytoca*) occurring within a neonatal unit, declared after three apparently unrelated colonized cases were identified. The admissions did not overlap in time and preliminary examination failed to reveal any likely cross contamination events. We explored the utility of whole genome sequencing (WGS) in conjunction with a standard outbreak investigation to help resolve the source of the outbreak and guide interventions to successfully halt transmission.

## Material and methods

### Setting

Caboolture Base Hospital (CBH) is a 280 bed regional hospital in South East Queensland, Australia. The Special Care Nursery (SCN) provides care for neonates greater than 32 weeks gestation or a birth weight of ≥1500g. The unit was designed to accommodate 12 neonates. However, due to demand is regularly forced to operate up to 14 beds. Active surveillance (rectal swabs) for multi-resistant organisms (MROs) including methicillin resistant *Staphylococcus aureus* (MRSA), multi-resistant Gram-negative (MRGN) bacteria and vancomycin resistant *Enterococcus* (VRE) is performed weekly and on all transfers in to the unit. All cases of MRO colonization or infection are reviewed by the infection control (IC) team and contact isolation is employed for all MRSA, VRE and MRGN (except ESBL-producing *Escherichia coli)*.

### Definitions, data collection and infection control responses

After declaration of the outbreak, enhanced surveillance was introduced with rectal swabbing of all SCN patients every 48 hours. All neonates with positive screening swabs were isolated in single rooms (when available) and with contact precautions (as per routine policy). Mothers of colonized babies were also screened. Direct observation of IC procedures and hand hygiene performance was performed by independent Hand Hygiene Australia (HHA) accredited IC staff. SCN procedures and policy compliance were reviewed.

### Microbial Surveillance

Rectal swabs were collected and plated onto selective chromogenic media (chromID™ ESBL agar; bioMérieux) and examined after 18-24 hours incubation in O_2_. Colonies which grew on selective media were identified by matrix-associated laser desorption/ionization-time-of-flight mass spectrometry (MALDI-TOF) (Vitek MS, bioMérieux) and antibiotic susceptibility determined using the Vitek2 system (bioMérieux). ESBL production was confirmed using combination disk diffusion, where an increase in the zone of inhibition ≥5mm by the addition of clavulanic acid (10 μg) to Cefotaxime (30 μg) or Ceftazidime (30 μg) disks suggests ESBL production.(23) After the incident case (case 1) was identified, any ceftriaxone-resistant *K. oxytoca* from neonatal cultures of any body-site were stored, and following the declaration of an outbreak (case 3) all stored and prospective isolates were sent for WGS. For comparison, additional isolates were sequenced, including ESBL-producing *E. coli* and ceftriaxone-susceptible *K. oxytoca* from neonates, as well as any *K. oxytoca* identified from maternal samples. An outbreak case was defined as any individual with *K. oxytoca* cultured from any site, which was determined to be resistant to ceftriaxone by Vitek2 (irrespective of ESBL phenotypic confirmatory testing).

### Environmental screening

Swabs were collected from sinks, drains, aerators of faucets, soap dispensers, damp surfaces, high use equipment and the humidification apparatus of humidicribs; and samples of consumables such as shampoo, paraffin and ultrasound gel were collected and plated onto selective chromogenic media and processed in the same way as the patient screening samples (see Microbial surveillance methods).

### Whole Genome Sequencing

All *K. oxytoca* isolates, from neonates, their mothers or the environment were submitted for WGS at the Queensland Health Forensic Scientific Services (FSS) laboratory in 5 batches between January 29^th^ 2018 and March 23^rd^ 2018. Genomic DNA was extracted using QIAamp DNA mini kits (Qiagen, Australia) and quantified by spectrophotometry (NanoDrop; ThermoFisher) and fluorometry (Quant-iT; ThermoFisher). Paired-end DNA libraries were prepared using Nextera XT kits (Illumina; Australia) and WGS was performed using Illumina NextSeq (150 bp paired end).

Using Nanopore sequencing (MinION; Oxford Nanopore Technologies) we assembled the complete genome of strain M82255 for reference. In brief, 1.5 µg of DNA was used as input for the 1D sequencing by ligation kit (SQK-LSK108) as per manufacturer’s instructions. The final library was loaded onto a FLO-MIN106 R9.4.1 flow cell and run for approximately 40 hours on a MinION.

### *In silico* multilocus sequencing typing, single nucleotide polymorphism (SNP) typing and resistance gene detection

Sample analysis was undertaken using the custom Queensland Genomics Health Alliance (QGHA) infectious diseases genomic analysis pipeline (version: dev-0.4.0). In brief, raw Illumina sequencing data was quality trimmed using Trimmomatic (version 0.36).(24) *In silico* sequence typing (ST) was preformed using SRST2 and the *K. oxytoca* typing scheme available from PubMLST (https://pubmlst.org/). Single nucleotide polymorphism (SNP) profiling and determination of core genome SNPs was undertaken by aligning the trimmed Illumina sequencing reads for each sample to the complete genome of M82255 using Nesoni (version 0.132) (https://github.com/Victorian-Bioinformatics-Consortium/nesoni). Resistance gene profiling was performed by screening the trimmed sequence reads for each isolate against the ARG-ANNOT resistance gene database using SRST2 (25, 26). Only genes with a minimum of 90% nucleotide sequence identity and 90% sequence coverage were reported.

### Phylogenetic analysis

A list of polymorphic positions conserved in all strains was established using Nesoni nway. Polymorphic substitution sites were concatenated to produce an alignment that was used to reconstruct the phylogeny. A recombination filtered, Maximum Likelihood phylogenetic tree was estimated using Gubbins version 2.2.0 (http://sanger-pathogens.github.io/gubbins/)(27) for the SNP alignments under the generalized time reversible (GTR) nucleotide substitution model with gamma correction for among site rate variation (ASRV).

### Accession numbers

Genome data has been deposited to NCBI under Bioproject PRJNA512395. Raw Nanopore and Illumina sequence read data has been deposited to the Sequence Read Archive (accessions: SRR89420303-SRR8420330). The complete genome of M82255 has been deposited to GenBank (accessions: CP035214-CP035216)

### Ethics

WGS activities for pathogen surveillance and infection control are covered by the Queensland Genomics Health Alliance (QGHA) clinical demonstration project (HREC/17/QFSS/6), with approval for waiver of consent.

## Results

### Species identification

Initial species characterization, using MALDI-TOF, identified strain M82255 as *Klebsiella oxytoca.* Taxonomic assignment was confirmed *in silico* using Kraken (28) by comparison of sequence read data for M82255 against the NCBI RefSeq database containing ∼25,000 complete bacterial, archaeal and viral genomes. However, after submission of the complete genome of M82255 to GenBank, a quality control test of the taxonomic assignment using average nucleotide identity (ANI) (29) revealed the genome of M82255 to be 98.903% identical to *K. michiganensis* (91.1% genome coverage) but only 92.455% similar to *K. oxytoca*. Further biochemical tests were performed on M82255 to identify its position within the genus *Klebsiella* (Supplementary table S1). Similar to the expected biochemical profile of *K. michiganensis*, M82255 was unable to produce urease (2) confirming the results of the ANI approach. Consequently, M82255 was reclassified as *K. michiganensis*.

### Incident cases

From the incident case to identification of the final case 188 days later, there were ten neonates colonized with *K. michiganensis* (Figure 1). One neonate was simultaneously co-colonized with ESBL-producing *E. coli*. During this period there were approximately 880 births and 163 admissions to the special care nursery (SCN). There were no cases of clinical infection caused by *K. michiganensis*. The average length of stay in SCN was 15.7 (range 5-35) days and there was a mean of 7.9 (range 2-15) days from admission to SCN to acquisition of ESBL-*K. michiganensis*. One patient neonatal sample was obtained from another hospital facility following discharge.

**Figure 1:**
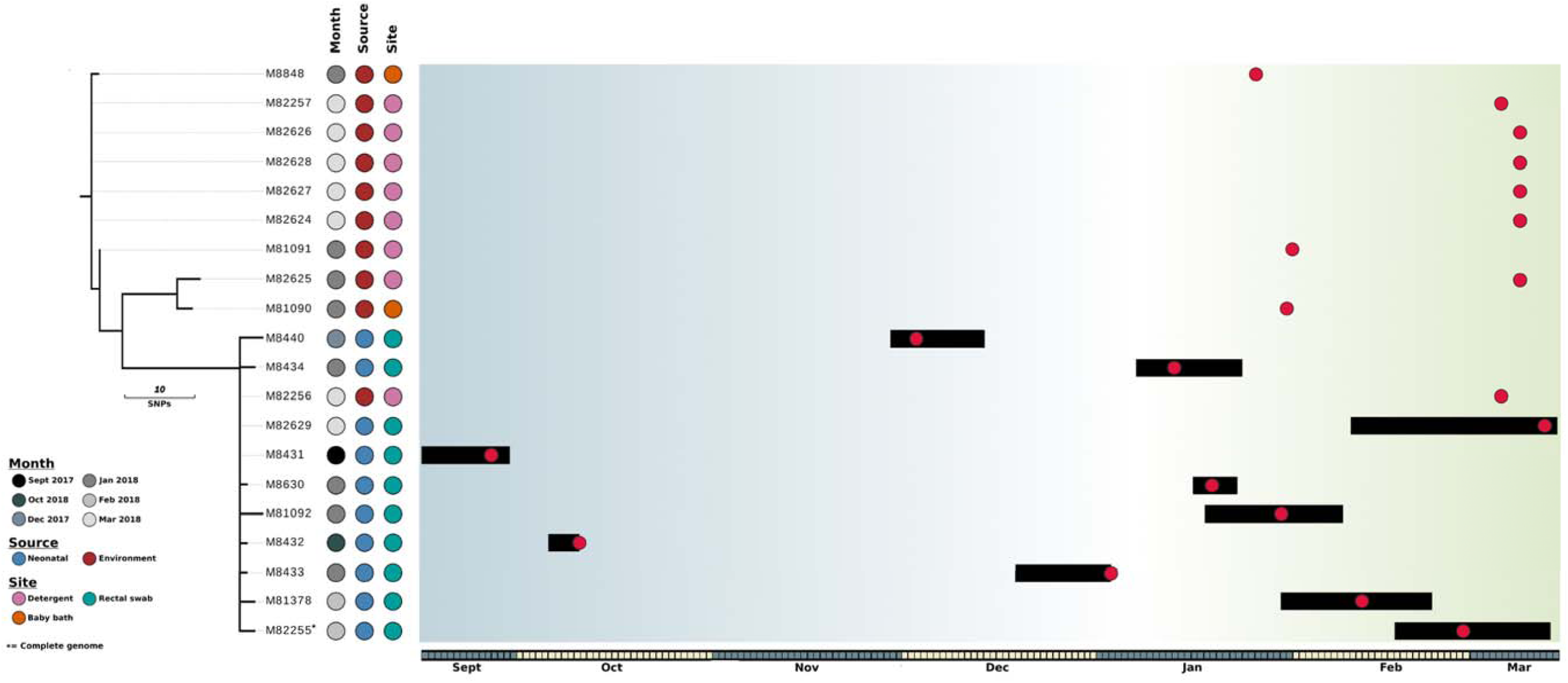
The evolutionary relationship of *K. michiganensis* isolates. Maximum likelihood phylogenetic tree built using 38 core genome SNPs relative to *K. michiganensis* strain M82255. The month of isolation, source (neonatal or environmental) and specific site of isolation are indicated as per the legend. Branches (black lines) represent the genetic distance between isolates in terms of SNPs. The scale is indicated. The outbreak timeline is presented adjacent to the tree. The timescale is represented on the bottom with alternate months colored blue and yellow. Each box in the timeline represent a period of 1 day starting at the 15th of September 2017. Back bars represent the admission periods for colonized patients and red dots represent dates on which the samples were isolated (including environmental isolates). Grey shading represent periods where there was no overlap in patient stays and green shading represents periods where multiple colonized patients were present in the SCN at the same time (yellow).

**Figure 2:**
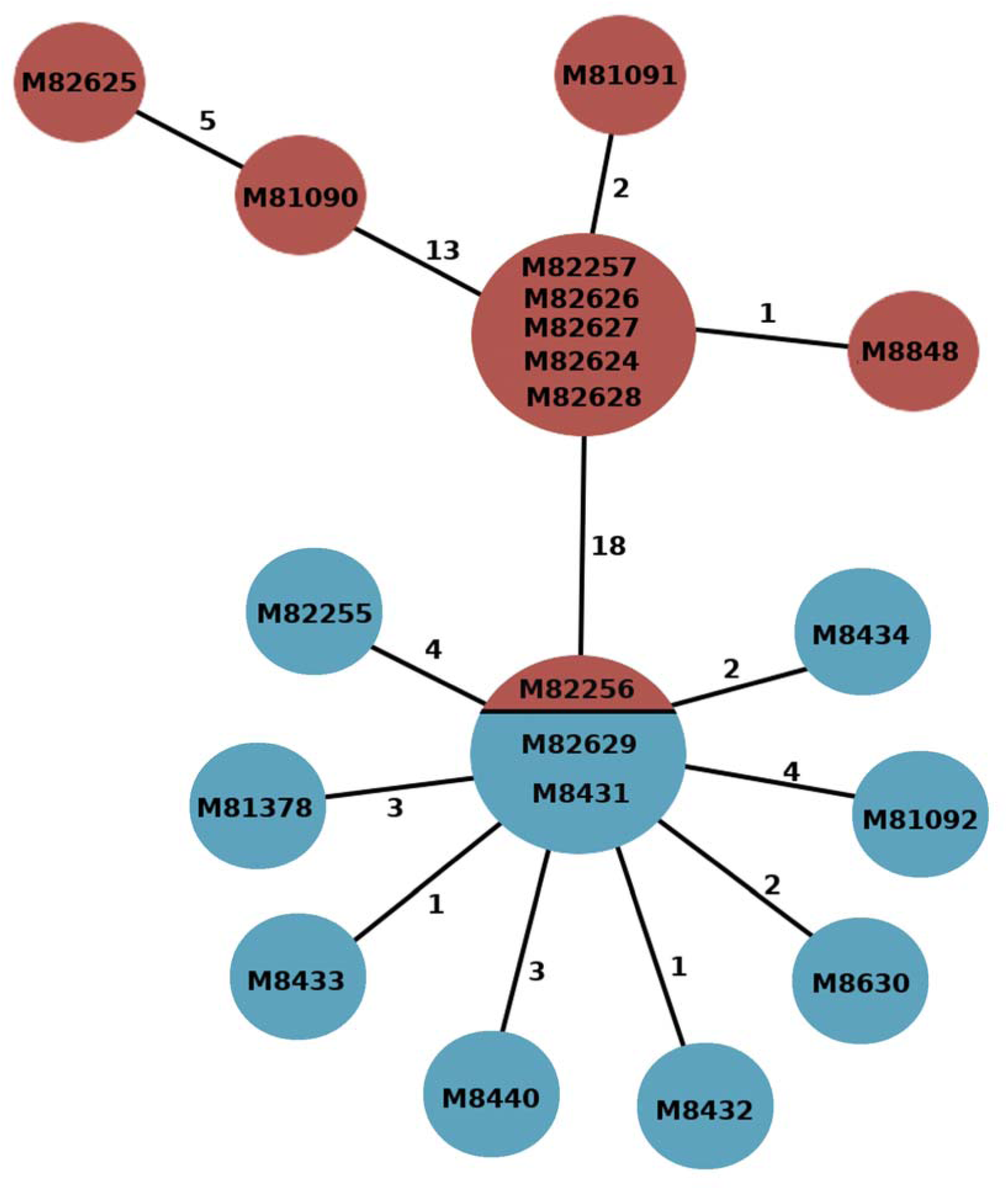
The evolutionary relationship of *K. michiganensis* strains. Minimal Spanning tree showing relationships between environmental (red) and patient (blue) samples. Numbers attached to branches represent number of SNPs between genomes. Isolate codes are represented within each node (if >1 code listed, then these were identical at the core genome level).

### Infection Control Observation

Direct observation of the SCN infection control (IC) procedures revealed that all staff hand hygiene (HH) performance was above the national benchmark (80%). However, significant opportunities for cross contamination (breakdown in HH, leaning on equipment, failure to observe contact isolation procedures) were noted in relation to non-staff (families, friends and SCN occupants). Additionally, incomplete compliance with microbial surveillance cultures on admission was noted at the start of the outbreak.

### Environmental Screening

Exhaustive screening of the SCN unit on multiple occasions failed to detect a source of environmental contamination. Environmental sampling was then extended to the maternity ward and birth suite. ESBL-producing *K. michiganensis* was identified from a baby-bath drain in a multi-purpose utility room containing laundry equipment (washing machine and laundry dryer), and a pair of baby baths (no longer used for bathing). However, an epidemiological link between the multi-purpose room in maternity and the colonized cases in SCN was not evident.

During IC unit observation it was noted that the hospital’s volunteer flower service used the multi-purpose room for cleaning flower vases. The flower services equipment was then sampled and a second *K. michiganensis* environmental isolate was retrieved from a sample of the flower service’s dishwashing detergent. The hospital’s detergent (Cleantec Emmy; Ecolab, Australia) was bought as bulk concentrate and then decanted into reusable detergent bottles for use around the hospital. Each drum of concentrate lasted approximately one month. *K. michiganensis* was not identified in the bulk concentrate at the time of testing, however sampling of detergent from around the hospital identified *K. michiganensis* from 7 of 13 bottles.

### Whole genome sequencing

A total of 21 organisms (10 patient-derived isolates, from 10 individuals and 11 environmental isolates) were included in the genomic analysis (Table 1). A single non-ESBL producing *K. oxytoca* (M8843) was identified from a maternal rectal swab and found to belong to the ST50 lineage and was excluded from further study and not considered part of the outbreak. All *K. michiganensis* strains possessed chromosomal OXY-type β-lactamases, as well as SHV-2 ESBL (in addition to LEN and TEM-1B β-lactamases). Resistance genes for aminoglycosides (*aac3-IId*, *aph(3’)-Ia*, *strA/B*) and sulphonamides (*sul1*) were also detected.

**Table 1:** Sample details

| **ID** | **SOURCE** | **GENDER** | **DATE SAMPLE** | **SAMPLE** | **ORGANISM** | **Hospital** |
| --- | --- | --- | --- | --- | --- | --- |
| **M8431** | Neonatal | Male | September-2017 | Rectal swab | *Klebsiella michiganensis* | Caboolture hospital |
| **M8432** | Neonatal | Female | October-2017 | Rectal swab | *Klebsiella michiganensis* | Caboolture hospital |
| **M8440** | Neonatal | Male | December-2017 | Skin swab | *Klebsiella michiganensis* | External hospital |
| **M8433** | Neonatal | Female | January-2018 | Rectal swab | *Klebsiella michiganensis* | Caboolture hospital |
| **M8434** | Neonatal | Female | January-2018 | Rectal swab | *Klebsiella michiganensis* | Caboolture hospital |
| **M8630** | Neonatal | Female | January-2018 | Rectal swab | *Klebsiella michiganensis* | Caboolture hospital |
| **M81092** | Neonatal | Female | January-2018 | Rectal swab | *Klebsiella michiganensis* | Caboolture hospital |
| **M81378** | Neonatal | Male | February-2018 | Rectal swab | *Klebsiella michiganensis* | Caboolture hospital |
| **M82255** | Neonatal | Female | February-2018 | Rectal swab | *Klebsiella michiganensis* | Caboolture hospital |
| **M82629** | Neonatal | Female | March-2018 | Rectal swab | *Klebsiella michiganensis* | Caboolture hospital |
| **M8848** | Environmental | - | January-2018 | Baby bath drain | *Klebsiella michiganensis* | Caboolture hospital |
| **M81090** | Environmental | - | January-2018 | Baby Bath | *Klebsiella michiganensis* | Caboolture hospital |
| **M81091** | Environmental | - | January-2018 | Detergent (Flower service) | *Klebsiella michiganensis* | Caboolture hospital |
| **M82256** | Environmental | - | March-2018 | Detergent (Milk Room) | *Klebsiella michiganensis* | Caboolture hospital |
| **M82257** | Environmental | - | March-2018 | Detergent (Nurses Station) | *Klebsiella michiganensis* | Caboolture hospital |
| **M82624** | Environmental | - | March-2018 | ICU Detergent Brush | *Klebsiella michiganensis* | Caboolture hospital |
| **M82625** | Environmental | - | March-2018 | Detergent (Pan Room) | *Klebsiella michiganensis* | Caboolture hospital |
| **M82626** | Environmental | - | March-2018 | Milk Room | *Klebsiella michiganensis* | Caboolture hospital |
| **M82627** | Environmental | - | March-2018 | Detergent (ICU) | *Klebsiella michiganensis* | Caboolture hospital |
| **M82628** | Environmental | - | March-2018 | Detergent (Pediatric ward) | *Klebsiella michiganensis* | Caboolture hospital |

To investigate the relationship between outbreak isolates at single nucleotide level reads from each isolate were mapped to the complete genome of *K. michiganensis* M82255. Phylogenetically, the *K. michiganensis* isolates formed two distinct clusters separated by 15 core genome SNPs (Figure 1; Supplementary table S2). Cluster 1 (SCN-C1 from this point on) was composed exclusively of environmental isolates (n=10). Cluster 2 (SCN-C2) was composed of the 10 patient isolates and a single environmental isolate, M82256.

WGS and an in-depth epidemiological investigation were used to determine if *K. michiganensis* transmission resulted from direct patient-to-patient transfer or through contact with a common environmental reservoir.

Between Jan 6^th^ and March 14^th^ (when the outbreak resolved) 6 colonized patients had overlapping stays in the SCN (Figure 1), four of whom were resident at the same time providing opportunity for isolate transmission between neonates. However, when WGS was used to establish relationships between these isolates there was no genetic evidence to support patient-to-patient transmission. SCN-C2 is composed of a tight cluster of closely related isolates (mean pairwise SNP distance of 3.3 SNPs).

Central to this cluster is a group of 3 isolates indistinguishable at the core genome SNP level. The cluster was composed of two isolates (M8431 and M82629) isolated from case 2 and case 10 respectively (hospital admissions separated by 147 days), and a single environmental isolate (M82256) collected from a detergent located in the SCN milk room. The remaining 8 patient isolates diverge directly from this central group but possess no common discriminatory SNPs, suggesting that each colonized patient independently acquired *K. michiganensis* from a common source, in all probability the SCN milk room detergent.

Within SCN-C1, environmental isolates collected from a baby bath drain (M8848) and detergent samples from 4 of the 8 tested detergent dispensing bottles (M82257, M82624, M82627 and M82628) were indistinguishable at the core genome level. The relatedness of isolates collected from the detergent suggests that contamination from a central source was likely responsible for introducing *K. michiganensis* into the hospital, possibly during dispensing of the bulk detergent or from the drum itself. Testing of the bulk detergent drum did not identify any biological contaminants. However, the high number of SNPs (14 core genome SNPs) separating environmental and patient isolates suggests that the outbreak strain may have been present in the local environment for some time.

### Outbreak Management

All identified cases were isolated under contact precautions. After isolation of *K. michiganensis* in the multi-purpose room, it was immediately decommissioned and refurbished as a single use facility. Redundant equipment (baby-baths, laundry equipment) was removed and the flower service was temporarily ceased. Reusable detergent bottles from across the hospital were destroyed and the detergent supply switched to pre-filled single use detergent bottles.

As the contamination of the detergent bottles may have originated from contaminated concentrate, a search for similar cases of ESBL-producing *K. michiganensis* (or *K. oxytoca*) at other hospitals served by Pathology Queensland was performed using the statewide laboratory information system, however there was no indication of unrecognized outbreaks at other facilities.

## Discussion

*Klebsiella* species are frequently implicated in outbreaks in the neonatal intensive care setting, although sources are not always evident despite extensive investigation and screening (30). For example, outbreaks of multi-drug resistant *K. oxytoca* have been previously linked to environmental reservoirs, with sources described as varied as handwashing sinks (7), wastewater drainage systems (9), water humidifiers for neonatal incubators (16), transducers used for blood pressure monitoring (31), and washing machines.(11) Contaminated medical solutions such as heparin (32), sodium chloride (33) or insulin (8) have also been described as a source.

Nosocomial outbreaks of *K. michiganensis* have not been reported previously. However, based on the average nucleotide identity (ANI) approach (34), numerous strains previously identified as *K. oxytoca* have now been reclassified *K. michiganensis*, including a carbapenem resistant strain, E718 (GenBank accession: CP003683), isolated from the abdomen of a renal transplant patient (35). In fact, of the seven complete *K. michiganensis* genomes on GenBank, including M82255 from the outbreak described here, all were originally classified as *K. oxytoca*. Consequently, it is possible that hospital outbreaks of *K. michiganensis* may go undetected as they are prone to misclassification.

Contaminated disinfectant has previously been recognized as a potential source of *K. oxytoca* sepsis in hospitalized infants.(36) Organism factors such as a mucoid phenotype via enhanced capsule formation may enable the bacteria to survive in this environment.(36) Additionally, it has been suggested that in *Klebsiella*, outer-membrane proteins, such as peptidoglycan-associated lipoprotein (Pal) and murein lipoprotein (LppA), contribute to resistance against detergents through modulating the integrity and selective impermeability of the cell membrane independently of LPS and capsule.(37) Strains of *K. oxytoca* have been described which are able to withstand active ingredients of commonly used detergents, such as sodium dodecyl sulfate (SDS).(38)

This report clearly demonstrates the effectiveness of high-throughput WGS as a tool to enhance and support established infection control outbreak responses. The high resolution available with WGS to compare outbreak strains was sufficient to rule out patient-to-patient transmission and suggested that an environmental reservoir might be the source of infection. The ensuing IC investigation successfully identified several environmental sites contaminated with *K. michiganensis.* WGS of the environmental isolates linked them, unequivocally, to patient isolates and identified a liquid detergent bottle in the milk dispensing room as the most likely source for onward transmission in the SCN. Washing detergent in the milk room was used by mothers to clean milk-expressing equipment resulting in cross-contamination between this equipment and expressed milk, which resulted in neonate colonization. Although the initial source of the contamination could not be confirmed, *K. michiganensis* isolates related to the outbreak were identified in liquid detergent dispensers located at different sites throughout the hospital. Liquid detergent dispensers are refilled from a central bulk drum of detergent concentrate. However, at the time of the outbreak the bulk concentrate was negative for *K. michiganensis.* It is possible that either a contaminated drum was introduced into the hospital, its contents dispensed and the drum disposed of prior to the outbreak or contamination occurred.

The hospital procedures and practice for preparation of cleaning materials were reviewed. It became apparent that detergent bottles would often be “topped-up” when approaching empty, rather than being emptied, washed out and left to dry before refilling. This practice would have allowed propagation of the organism in the detergent bottles after contamination. In response to this re-usable detergent bottles around the hospital were collected and destroyed and single use pre-diluted detergent bottled were sourced. Although the topping up of detergent bottles was non-compliant with existing procedures, from the point of view of the maintenance staff is was intuitive and convenient. This gap between “practice imagined” and “practice performed” illustrates the importance of direct observation of staff practice and procedures when investigating an outbreak, the importance of all hospital staff to understand the infection control policies relevant to them and for a well-resourced, active hospital infection control team.

## Conclusions

Environmental reservoirs are increasingly recognized as a source of outbreaks involving MDR pathogens. We demonstrate how whole genome sequencing, in conjunction with standard epidemiological investigation, was instrumental in elucidating the source of ESBL-producing *K. michiganensis* in a neonatal unit and directing effective measures to interrupt transmission. Liquid detergent can become contaminated with pathogenic bacteria and should be considered as a potential source in such outbreaks.

## Acknowledgements

We thank the scientists at Pathology Queensland for their assistance in the laboratory work.

