## Supplementary table S1 for "Genomic investigation reveals contaminated detergent as the source of an ESBL-producing *Klebsiella michiganensis* outbreak in a neonatal unit"

**Table S1:** Phenotypic biochemical characteristics of *K. oxytoca* and *K. michiganesis*

|  | ***K. oxytoca* (ATCC 131182)** | ***K. michiganesis*** | **M82255** |
| --- | --- | --- | --- |
| **Indole production** | + | + | + |
| **Urease production** | + | - | - |
| **Voges-Proskauer** | + | + | + |
| **ONPG** | + | + | + |
| **Ornithine decarboxylation** | - | - | - |
| **Motility** | - | - | - |
| **Catalase** | + | + | + |
| **Adonitol** | + | + | + |
| **Glucose** | + | + | + |
| **Mannitol** | + | + | + |
| **Inositol** | + | + | + |
| **Sucrose** | + | + | + |
| **Arabinose** | + | + | + |
| **Rhamnose** | + | + | + |
| **Melibiose** | + | + | + |
| **Sorbitol** | + | + | + |
| **H2S** | - | - | - |
